# Dynamic Regulation of Atg4 Protease and Autophagy by Dithiothreitol and Iron in *Alternaria alternata*

**DOI:** 10.64898/2026.05.04.722700

**Authors:** Hsin-Yu Lu, Celine Yen Ling Choo, Pei-Ching Wu, Kuang-Ren Chung

## Abstract

Autophagy is a critical cellular process regulated by Atg proteins, yet its modulation by redox-active compounds and iron remains incompletely understood. Here, we investigated the effects of dithiothreitol (DTT) and iron on autophagy and on AaAtg4 protease activity in the plant-pathogenic fungus *Alternaria alternata*. Using GFP-tagged AaAtg8, fluorescence microscopy and proteolysis assays revealed that DTT markedly enhanced autophagic vacuole formation and GFP release, indicating increased autophagic flux. Western blot analyses confirmed that DTT promoted AaAtg8 lipidation, while co-treatment with hydrogen peroxide (H₂O₂) suppressed this modification. AaAtg4 was constitutively active and could process AaAtg8 regardless of DTT supplementation, whereas moderate DTT concentrations elevated AaAtg4 protein abundance and phosphorylation. Bimolecular fluorescence complementation assays demonstrated that DTT, but not iron, facilitated AaAtg4–AaAtg8 interactions and vacuolar localization, whereas H₂O₂ counteracted these effects. Notably, combined DTT and H₂O₂ sustained autophagy at a low but stable level, suggesting a redox balance in autophagic regulation. Iron supplementation selectively destabilized AaAtg8 and modulated AaAtg4 phosphorylation in a concentration-dependent manner, without altering autophagy or protease activity. Collectively, these findings demonstrate that DTT enhances autophagy primarily by promoting AaAtg8 lipidation, AaAtg4 phosphorylation, and AaAtg4–AaAtg8 complex formation, while exerting minimal influence on AaAtg4 protease activity. In contrast, ion regulates autophagy flux through its effects on AaAtg4 phosphorylation and AaAtg8 stability, without significantly altering AaAtg4 protease activity, AaAtg8 lipidation, or AaAtg4–AaAtg8 interactions. Together, this work underscores the intricate interplay between redox signaling, nutrient cues, and autophagy regulation in *A. alternata*.

**IMPORTANCE:** This study provides critical new insights into how redox-active compounds and iron modulate autophagy in the plant-pathogenic fungus *Alternaria alternata,* a pathogen of agricultural relevance. By dissecting the distinct roles of DTT, hydrogen peroxide, and iron in regulating AaAtg8 lipidation, AaAtg4 phosphorylation, and AaAtg4–AaAtg8 interactions, our findings reveal that autophagy is not simply a constitutive process but is finely tuned by redox balance and nutrient cues. This work advances the fundamental understanding of autophagy regulation in filamentous fungi, highlights the interplay between oxidative stress and protease activity, and establishes a framework for exploring how environmental factors shape fungal pathogenicity. Ultimately, these insights may inform novel strategies to mitigate crop fungal diseases by targeting autophagic pathways.

## INTRODUCTION

Autophagy is a conserved cellular degradation pathway that plays a critical role in maintaining homeostasis under stress conditions. Autophagy is controlled by more than 40 autophagy-related (Atg) proteins in the yeast *Saccharomyces cerevisiae* (1, 2). Among them, the ubiquitin-like protein Atg8, which is required for phagophore elongation and autophagosomal maturation, plays a central role in autophagy. The glycine residue at the C-terminus of Atg8 is covalently linked to phosphatidylethanolamine (PE) to form an Atg8-PE conjugate, a process called lipidation. Lipidation and subsequent localization to autophagic membranes are tightly regulated by the cysteine protease Atg4 (3, 4). Atg8-PE is attached to autophagosome membranes. At the later stage of autophagy, Atg4 also functions to remove Atg8 from autophagosome membranes (a process called delipidation) to reimplement the cytosolic Atg8 pool (5, 6). Proper coordination between Atg8 processing and Atg4 activity ensures efficient autophagosome formation and turnover.

Redox signals, particularly those mediated by reducing agents and reactive oxygen species (ROS), are important modulators of autophagy (7, 8). Oxidative stress can induce autophagy and affect phagophore expansion, autophagosome maturation, cargo delivery, degradation, and recycling, as well as autophagy-related gene expression (9). At low concentrations, ROS function as signaling messengers, modulating autophagy by oxidizing Atg4 and other autophagy-related proteins (7). Moderate ROS levels can further stimulate autophagy to eliminate oxidized proteins and damaged organelles. In contrast, excessive ROS causes irreversible oxidative damage, impairs autophagy, and may lead to autophagic cell death or ferroptosis (10). In yeast, Atg4 activity is tightly controlled by the intracellular redox environment: reversible oxidation of cysteine residues forms disulfide bonds that inactivate Atg4, while reducing conditions restore its activity, enabling the removal of glycine from Atg8 and the deconjugation of Atg8-PE (6,11). Importantly, ROS-induced oxidation of Atg4 can be reversed by reducing agents such as dithiothreitol (DTT) or thioredoxin (11, 12). This dynamic cycle of activation and inactivation ensures precise regulation of autophagosome biogenesis and autophagic flux. DTT is known to alter protein folding and disulfide bond formation (13); however, its impact on autophagy-related proteins remains uncharacterized in phytopathogenic fungi, including *Alternaria alternata*.

The phytopathogenic fungus *A. alternata* can resist high levels of ROS partly by mitigating ROS stress via autophagy-mediated peroxisome degradation, a process called pexophagy (14). The *A. alternata* Atg8-deficient mutant (Δ*AaAtg8*) fails to efficiently degrade peroxisomes, leading to peroxisome accumulation, hypersensitivity to oxidative stress, and virulence reduction (15). Hydrogen peroxide (H_2_O_2_) triggers autophagy formation and the translocation of peroxisomes into vacuoles. In *A. alternata*, AaAtg8 physically interacts with AaAtg4, whose functions are indispensable for AaAtg8 processing and autophagosome formation (16). AaAtg4 modification and its involvement in the autophagic process are modulated by redox status, consistent with findings in yeast (12). H_2_O_2_ differentially affects AaAtg4 and autophagy, depending on the concentration tested. H₂O₂ levels at 10 mM enhance AaAtg4 phosphorylation and autophagy, whereas excessive H₂O₂ suppresses both. H₂O₂ increases AaAtg8-PE formation in a threshold-dependent manner, peaking at 20 mM. However, H₂O₂ suppresses AaAtg4 protease activity and AaAtg4-AaAtg8 binding in a concentration-dependent manner. In contrast to yeast (17), AaAtg4 does not interact with the AaAtg1serine/threonine kinase, nor is it phosphorylated in *A. alternata* (16).

In addition to pexophagy, iron is required for mitigating ROS toxicity in *A. alternata*. When siderophore-mediated iron acquisition is impaired, the fungus exhibits heightened sensitivity to oxidative stress (18). The heightened sensitivity to H₂O₂ exhibited by the siderophore-biosynthetic mutant can be mitigated through the exogenous addition of ferric iron. Iron is an essential micronutrient that plays diverse roles in cellular physiology. Iron acts as a cofactor in enzymatic reactions, a regulator of oxidative balance, and a modulator of signaling pathways (19). In filamentous fungi, iron availability is tightly linked to growth, stress responses, and pathogenicity (20). Autophagy is involved in regulating cellular iron homeostasis and adaptation to nutrient fluctuations. When autophagy is impaired, *A. alternata* fails to synthesize siderophores effectively, resulting in increased sensitivity to iron and the iron chelator bathophenanthroline disulfonic acid (15, 21, 22). While the regulation of autophagy has been extensively studied in yeast and higher eukaryotes, the influence of iron on the stability and activity of Atg4 remains poorly understood.

In this study, we investigated the effects of DTT and iron on autophagy in *A. alternata*. Our findings revealed that DTT enhanced autophagy by promoting AaAtg8 lipidation and facilitating AaAtg4–AaAtg8 interactions, but did not directly alter AaAtg4 protease activity. Moreover, we showed that DTT counteracted the inhibitory effects of hydrogen peroxide (H₂O₂) on autophagy, sustaining basal autophagic activity under oxidative stress. Our results also demonstrated that iron modulated the stability of the AaAtg8 protein and autophagic flux without altering AaAtg4 proteolytic activity and that iron regulated AaAtg4 phosphorylation in a concentration-dependent manner. These results highlight an important role of redox balance in regulating autophagy and provide new insights into how reducing conditions modulate the Atg machinery in filamentous fungi.

## RESULTS

### DTT enhances autophagy but does not affect AaAtg4 protease activity

The WT/GFP-AaAtg8 strain, which expresses a synthetic GFP translationally fused to the N terminus of the AaAtg8 protein, was used to assess the effect of DTT on autophagy. Fluorescence microscopy showed that, after being shifted to potato dextrose broth (PDB), the strain exhibited weak green fluorescence evenly distributed throughout the hyphae (Fig. 1). In contrast, when shifted to PDB supplemented with low-to-moderate levels of DTT for 4 h, the fluorescence appeared as aggregated puncta that colocalized with vacuoles (stained with CMAC), indicating the formation of autophagic vacuoles (AV). Elevating the DTT concentration from 0.5 mM to 10 mM resulted in a proportional increase in green fluorescence intensity within the vacuoles, with the brightest and most sharply defined puncta observed at 10 mM. In contrast, no aggregated puncta were detected; instead, weak green fluorescence was uniformly distributed along the hyphae, excluding vacuoles, following treatment with 30 mM DTT.

**FIG 1.**
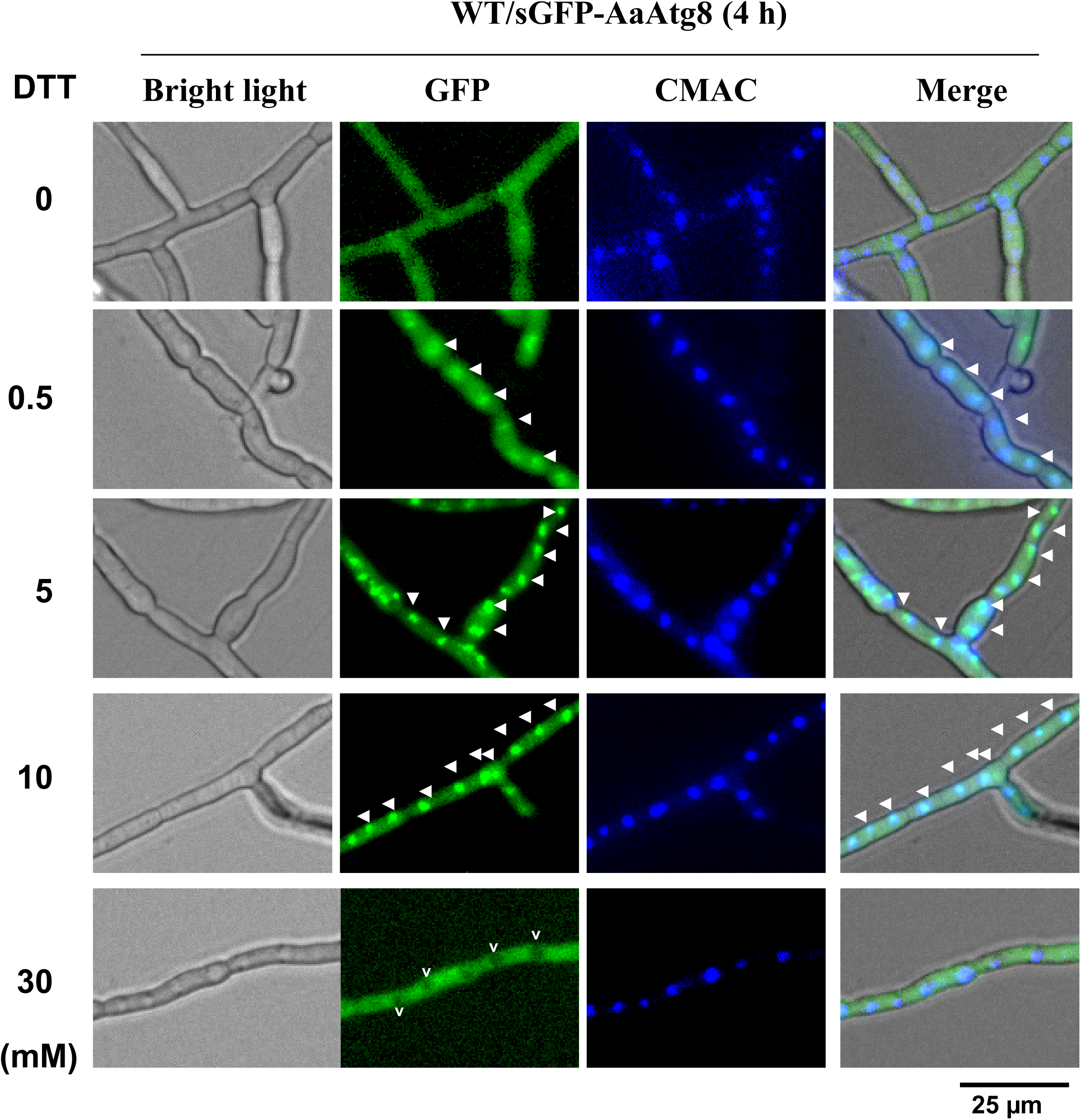
DTT enhances autophagy. Fluorescence microscopy of the WT/GFP-AaAtg8 strain revealed a diffuse GFP signal throughout the hyphae following transfer to PDB. In contrast, punctate GFP fluorescence colocalizing with vacuoles (V) is visible when the medium was supplemented with DTT, indicating the formation of autophagic vacuoles (highlighted by white arrowheads). Vacuoles were visualized using CMAC staining. The fungal strain was cultured in PDB for 24 h, after which the mycelium was harvested, washed, transferred to PDB containing DTT, incubated for an additional 4 h, and examined using a fluorescence-equipped microscope.

To further validate the role of DTT in enhancing autophagy, GFP-AaAtg8 proteolysis was assessed by measuring the release of free GFP during autophagy. Western blot analysis showed that only low levels of free GFP (∼8%) were detected in the wild type 6 h post incubation in PDB (Fig. 2A). Upon transfer to PDB supplemented with DTT at concentrations of 0.2, 5, 10, and 30 mM, the levels of free GFP increased to 16%, 60%, 61%, and 50%, respectively. To examine AaAtg4 protease activity, the WT/AaAtg8-GFP strain, expressing GFP fused to the C terminus of AaAtg8, was analyzed. Western blotting revealed complete conversion of AaAtg8-GFP to free GFP under all conditions tested, whether shifted to PDB alone or PDB supplemented with DTT (Fig. 2B). Based on free GFP band intensities, DTT treatment increased free GFP levels, with the highest observed at 5 mM compared to the control. No AaAtg8-GFP fusion protein was detected in any treatment.

**FIG 2.**
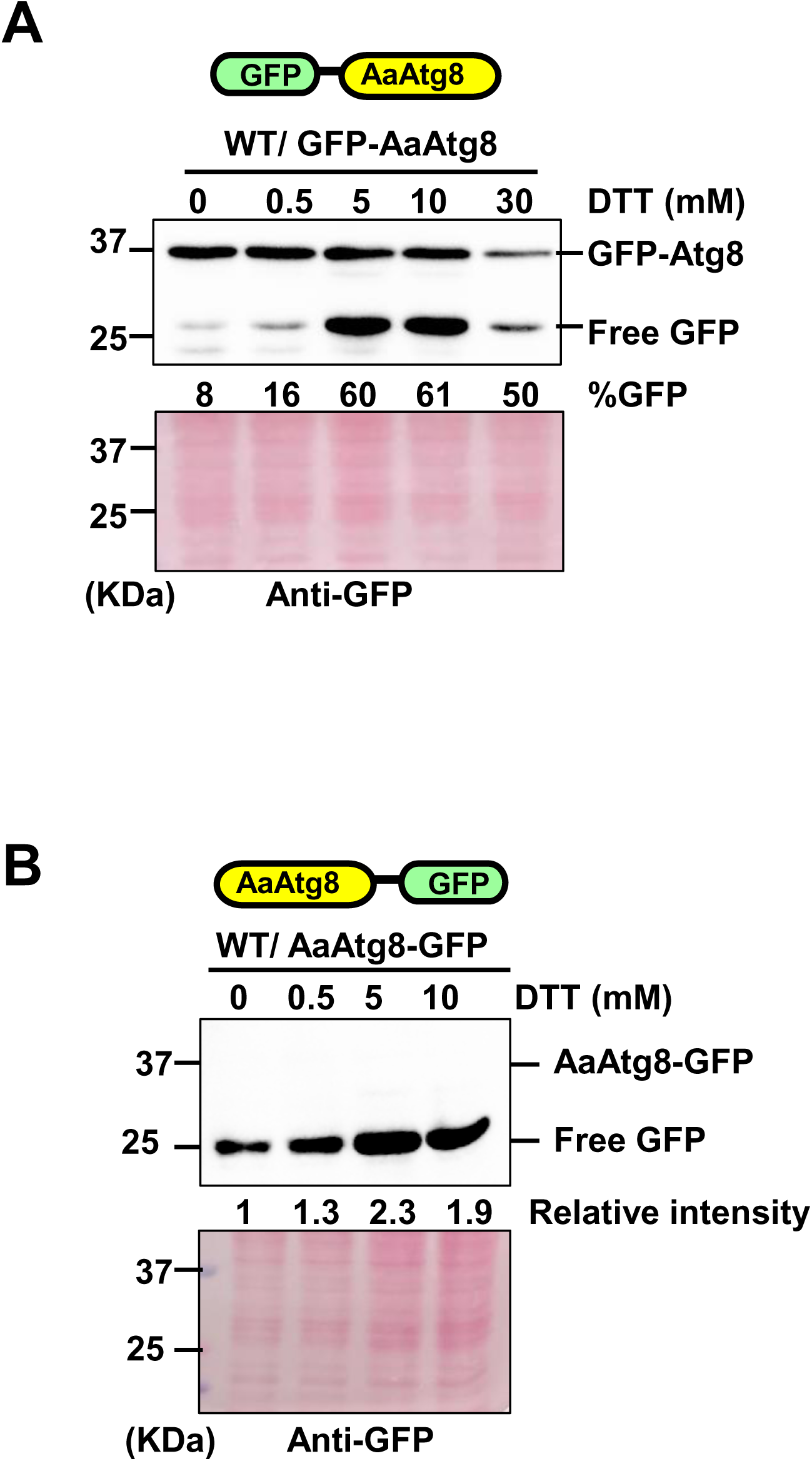
AaAtg8 proteolytic assays reveal that DTT is required for autophagy but does not influence AaAtg4 protease activity. (A) Western blot analysis of GFP-AaAtg8 proteolysis shows low levels of free GFP after 6 h in PDB, but increased release in cultures supplemented with DTT. The percentage of free GFP released from GFP–AaAtg8 was calculated by dividing the intensity of the free GFP band by the total intensity of free GFP plus GFP–AaAtg8, and then multiplying by 100%. (B) Western blot analysis of AaAtg4 protease activity in the WT/AaAtg8-GFP strain reveals complete conversion of AaAtg8-GFP to free GFP under all conditions, with band intensity enhanced by increasing DTT concentrations. No AaAtg8-GFP fusion protein is detected, although higher DTT concentrations increase free GFP band intensity. The band intensity of the mock control (without DTT) was set to 1, and the relative band intensity was calculated relative to the mock control.

### DTT facilitates the interaction between AaAtg4 and AaAtg8

The WT/GFP^N^-AaAtg8+GFP^C^-AaAtg4 (BiFC) strain, co-expressing AaAtg8 fused to the N-terminal half of GFP (GFP^N^) and AaAtg4 fused to the C-terminal half of GFP (GFP^C^), was used to examine the effect of DTT on the interaction between AaAtg8 and AaAtg4. Because GFP^N^ and GFP^C^ were truncated, green fluorescence was only produced when AaAtg8 physically interacts with AaAtg4. When cultured in PDB for 2 h, the strain exhibited weak green fluorescence in the hyphae (Fig. 3). The fluorescence intensity increased after 24 h of incubation. Notably, the signal became brighter when the strain was grown in PDB supplemented with 0.5 or 5 mM DTT, but diminished markedly when cultured in PDB containing 10 mM DTT.

**FIG 3.**
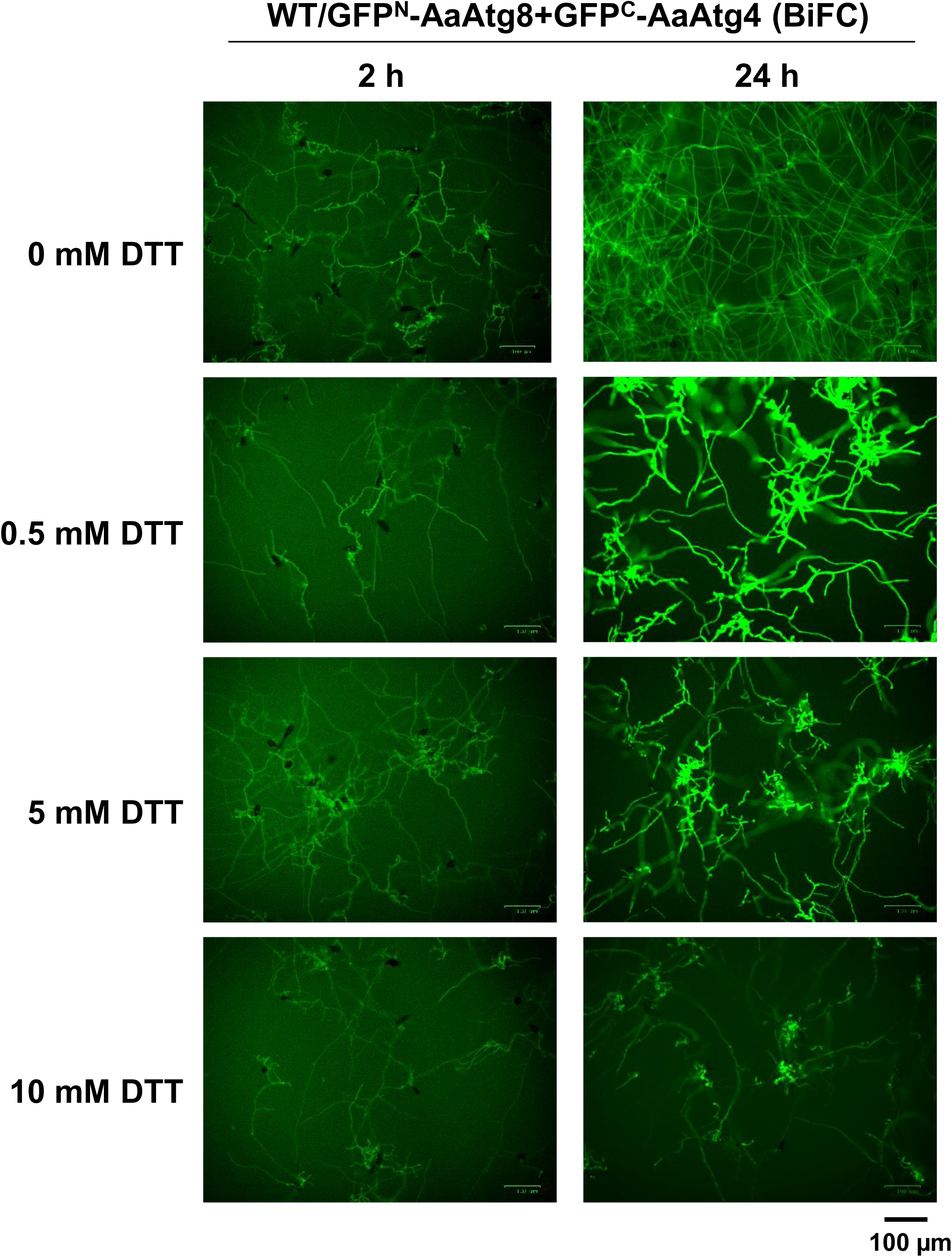
DTT facilitates the interaction between AaAtg4 and AaAtg8. BiFC analysis using the WT/GFPN-AaAtg8+GFPC-AaAtg4 strain demonstrates that AaAtg8 and AaAtg4 physically interact, as indicated by reconstituted GFP fluorescence. The hyphae show weak green fluorescence after 2 h in PDB, with intensity increasing after 24 h. Supplementation with 0.5 or 5 mM DTT further enhances fluorescence, indicating stronger AaAtg8–AaAtg4 interaction. In contrast, incubation with 10 mM DTT markedly reduces the signal, suggesting that excessive DTT impairs the interaction.

### DTT increases AaAtg8 lipidation, and H_2_O_2_ suppresses AaAtg8 lipidation induced by DTT

The WT/HA-AaAtg8 strain was used to investigate the effect of DTT on AaAtg8 lipidation, a process in which PE is conjugated to the C terminus of AaAtg8. Western blot analysis consistently detected an 18-kDa HA-AaAtg8 protein across all treatments. A 16-kDa band, corresponding to HA-AaAtg8-PE, was observed in strains incubated in PDB supplemented with 2, 5, 10, or 30 mM DTT, with the strongest signal detected at 10 mM DTT (Fig. 4A). Due to its increased hydrophobicity, HA-AaAtg8-PE migrated faster than HA-AaAtg8 in SDS-PAGE gels (23). Notably, HA-AaAtg8-PE was absent in samples from strains co-incubated with 10 mM DTT and varying concentrations of H₂O₂ (10, 20, or 30 mM) (Fig. 4B).

**FIG 4.**
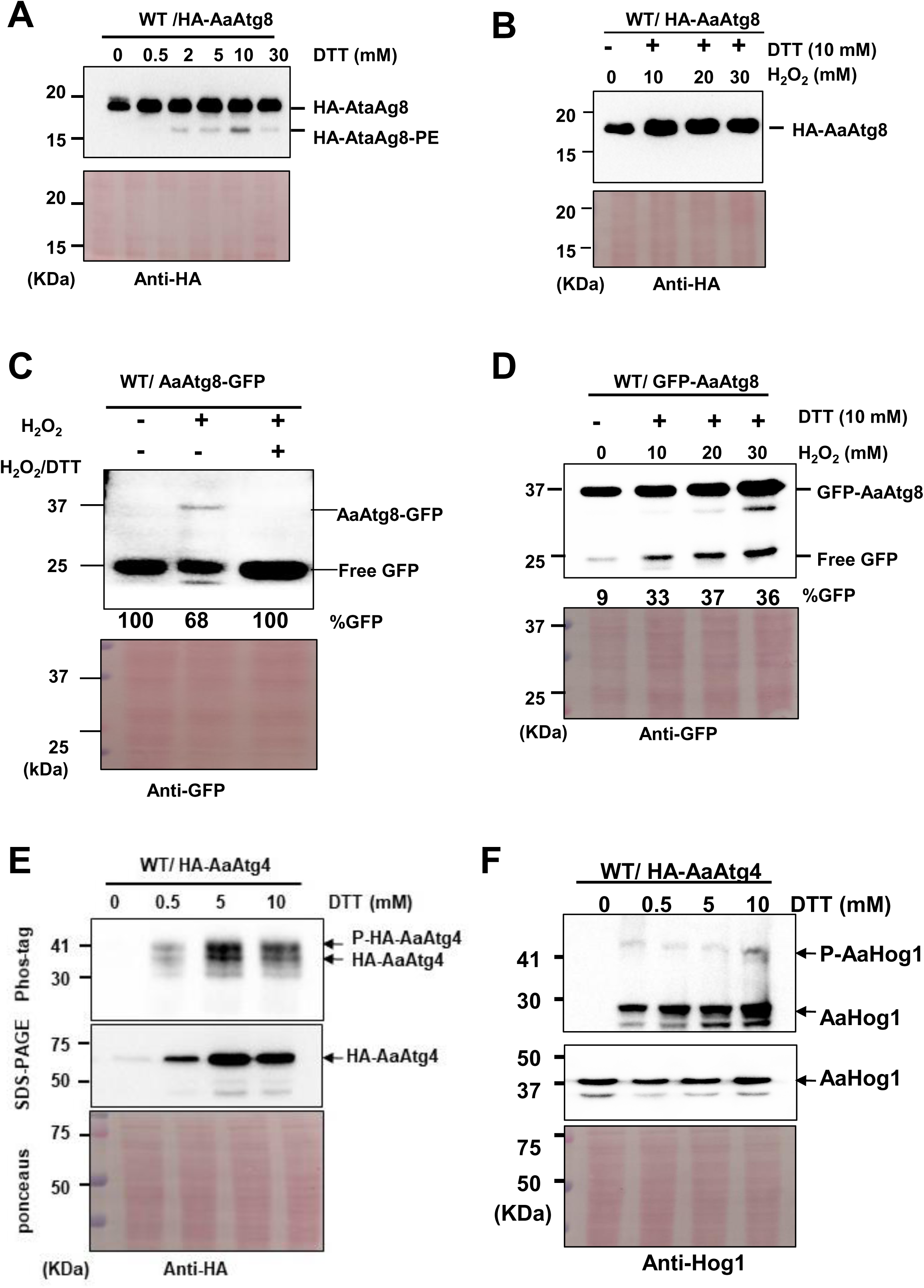
DTT promotes AaAtg8 lipidation and AaAtg4 phosphorylation, while H₂O₂ counteracts these effects and modulates autophagy. (A) Western blot analysis of WT/HA-AaAtg8 strains reveals that DTT induces AaAtg8 lipidation, as shown by the faster-migrating HA-AaAtg8-PE band. Lipidation is strongest at 10 mM DTT. (B) Co-incubation with H₂O₂ (10–30 mM) suppresses DTT-induced AaAtg8 lipidation, eliminating the HA-AaAtg8-PE band. (C) In WT/AaAtg8-sGFP strains, AaAtg4 protease activity is reduced by 30 mM H₂O₂ and abolished when both 30 mM H₂O₂ and 10 mM DTT are present, as indicated by loss of the AaAtg8-GFP band. Band intensities were measured by ImageJ. The percentage of free GFP released from GFP–AaAtg8 was calculated by dividing the intensity of the free GFP band by the total intensity of free GFP plus GFP–AaAtg8, and then multiplying by 100%. (D) GFP-AaAtg8 proteolysis assays show that autophagy is minimal in the absence of stressors but sustains at a moderate level (33–37% free GFP) when DTT and H₂O₂ co-exist. (E) WT/HA-AaAtg4 strains demonstrate increased AaAtg4 protein abundance and phosphorylation in response to DTT, peaking at 5 mM, but are reduced at 10 mM. (F) Phos-tag assays reveal that AaHog1 phosphorylation remains low and unaffected by DTT, consistent with its role in ROS signaling.

### DTT counteracts the H_2_O_2_ effect on AaAtg4 protease activity

The WT/AaAtg8-sGFP strain showed a strong AaAtg4 protease activity in PDB without DTT or H₂O₂ (Fig. 4C). Exposure to 30 mM H₂O₂ for 4 h slightly suppressed AaAtg4 activity, as indicated by the presence of a faint AaAtg8-GFP band. In contrast, the AaAtg8-GFP band was completely absent when both 30 mM H₂O₂ and 10 mM DTT were present, suggesting strong inhibition of AaAtg4 protease activity under these conditions.

### The co-existence of DTT and H_2_O_2_ sustains autophagy at a steady level

In the GFP-AaAtg8 proteolysis assay, only trace amounts of free GFP (∼9%) were detected in PDB when both H₂O₂ and DTT were absent (Fig. 4D), indicating minimal autophagic activity. However, the combination of 10 mM DTT with 10 mM H₂O₂ induced the release of approximately 33% free GFP. Notably, further increases in H₂O₂ concentration did not enhance GFP release. When 20 mM and 30 mM H₂O₂ were combined with 10 mM DTT, the levels of free GFP remained comparable, at 37% and 36%, respectively.

### DTT increases the AaAtg4 protein and phosphorylation levels

The WT/HA-AaAtg4 strain was employed to assess AaAtg4 phosphorylation in response to DTT. Western blot analysis showed that AaAtg4 protein was barely detectable when the strain was incubated in PDB; however, its levels increased upon transfer to PDB supplemented with DTT compared to PDB alone (Fig. 4E). AaAtg4 abundance increased progressively with DTT concentrations ranging from 0.5 mM to 10 mM. Consistently, phos-tag analysis demonstrated higher levels of AaAtg4 phosphorylation in PDB supplemented with DTT than in unsupplemented PDB. When the strain was shifted to PDB containing 10 mM DTT, both AaAtg4 protein accumulation and phosphorylation were similar to the levels observed with 5 mM DTT.

### DTT does not affect AaHog1 phosphorylation

Previous studies demonstrated that AaAtg4 physically interacts with AaHog1, a mitogen-activated protein kinase implicated in resistance to ROS and salt stress (24). Moreover, H₂O₂ has been shown to enhance AaHog1 phosphorylation in a dose-dependent manner (16). In contrast, phos-tag assays revealed that AaHog1 was phosphorylated at a low level regardless of the presence or absence of DTT (Fig. 4F).

### DTT promotes vacuolar localization of the AaAtg4-AaAtg8 complex, whereas H_2_O_2_ counteracts this effect

In the WT/sGFPN-AaAtg8+sGFPC-AaAtg4 strain, green fluorescence was observed in the hyphae but excluded from the vacuoles. Upon treatment with 0.1 mM DTT for 24 h, the strain exhibited uniform green fluorescence throughout the hyphae, including the vacuoles. However, when 40 mM H₂O₂ was added to the DTT-treated culture for 6 h, green fluorescence was no longer detected in the vacuoles (Fig. 5). Adding H₂O₂ at concentrations lower than 30 mM had no such effect (data not shown).

**FIG 5.**
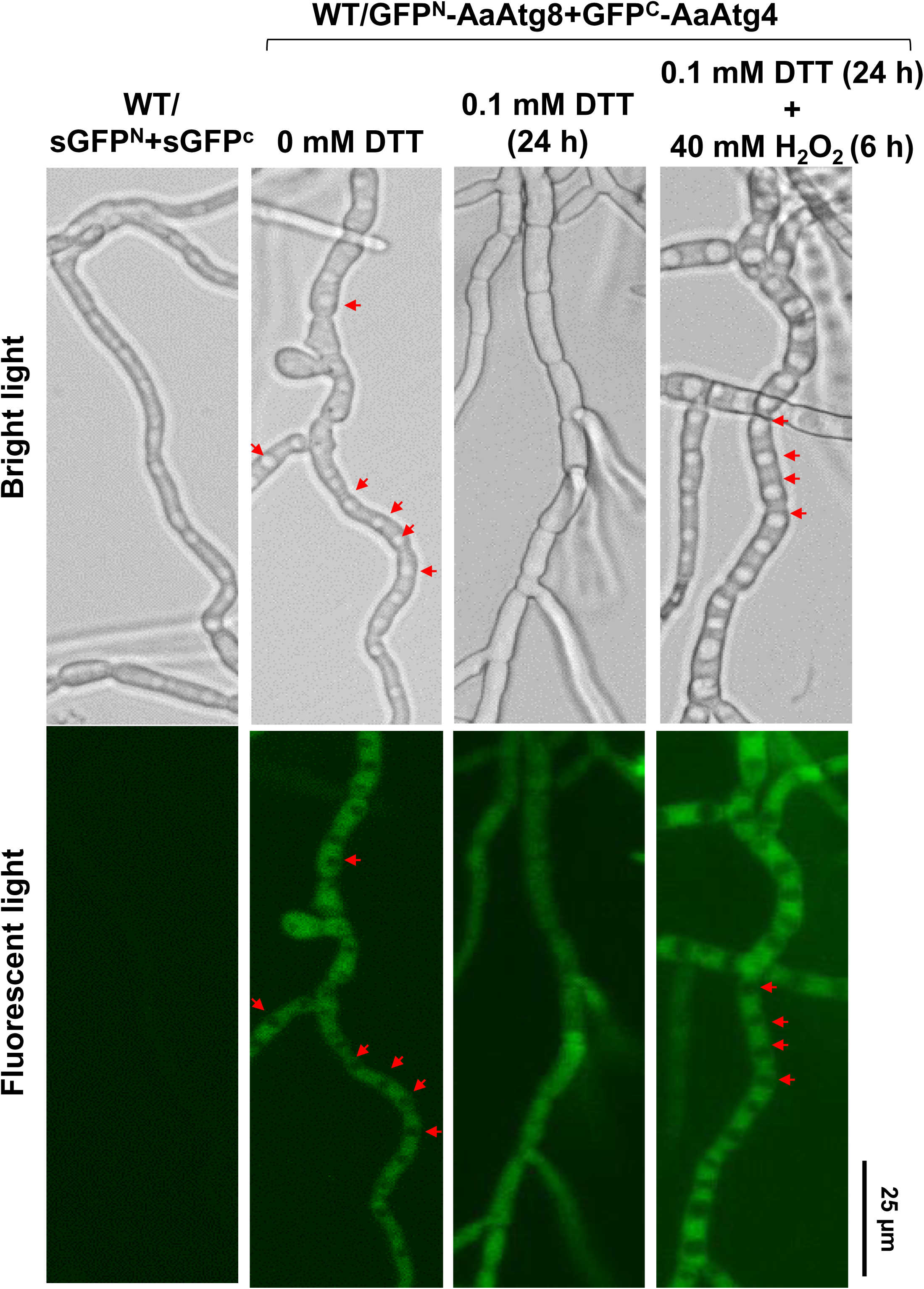
DTT promotes vacuolar localization of the AaAtg4–AaAtg8 complex, whereas H₂O₂ counteracts this effect. In the WT/sGFPN-AaAtg8+sGFPC-AaAtg4 strain, green fluorescence is observed in the hyphae but excluded from the vacuoles (indicated by red arrows). Treatment with 0.1 mM DTT for 24 h results in uniform green fluorescence throughout the hyphae, including the vacuoles. Subsequent addition of 40 mM H₂O₂ for 6 h abolishes vacuolar fluorescence, restricting the signal to the cytoplasm. Lower concentrations of H₂O₂ (<30 mM) do not affect vacuolar localization (data not shown).

### Iron modulates AaAtg8 stability without impacting autophagy or AaAtg4 activity

In the AaAtg8 proteolysis assay, the WT/GFP-AaAtg8 strain grown in minimal medium (MM) supplemented with ferric chloride (FeCl₃) or ferrous sulfate (FeSO₄) exhibited higher levels of free GFP compared to growth in MM alone (Fig. 6A). In contrast, treatment with bathophenanthrolinedisulfonic acid (BPS, an iron chelator) resulted in free GFP levels comparable to the unamended control. Deletion of the *AaAtg4* gene completely abolished the release of free GFP from GFP-AaAtg8 under all tested conditions. In the AaAtg4 protease assay, free GFP was fully released from AaAtg8-GFP in the wild type grown in MM or MM supplemented with FeCl_3_ or BPS (Fig. 6B), whereas no free GFP was detected in the Δ*AaAtg4* strain. Notably, GFP-AaAtg8 and AaAtg8-GFP fusion protein levels were markedly reduced following FeCl_3_ treatment compared to FeSO_4_ or other treatments. In the wild type, the release of free GFP from GFP-AaAtg8 increased as the concentration of FeCl₃ or FeSO₄ rose. However, GFP-AaAtg8 levels decreased progressively with increasing FeCl_3_ concentration (Fig. 6C), while FeSO_4_ had only a less effect (Fig. 6D). A similar FeCl_3_-dependent reduction of GFP-AaAtg8 levels was observed in the Δ*AaAtg4* strain (Fig. 6E). Conversely, GFP-AaAtg8 levels increased with higher concentrations of FeSO_4_ (Fig. 6F) or BPS (Fig. 6G) in the Δ*AaAtg4* strain. These trends were consistently reproduced across independent experiments.

**FIG 6.**
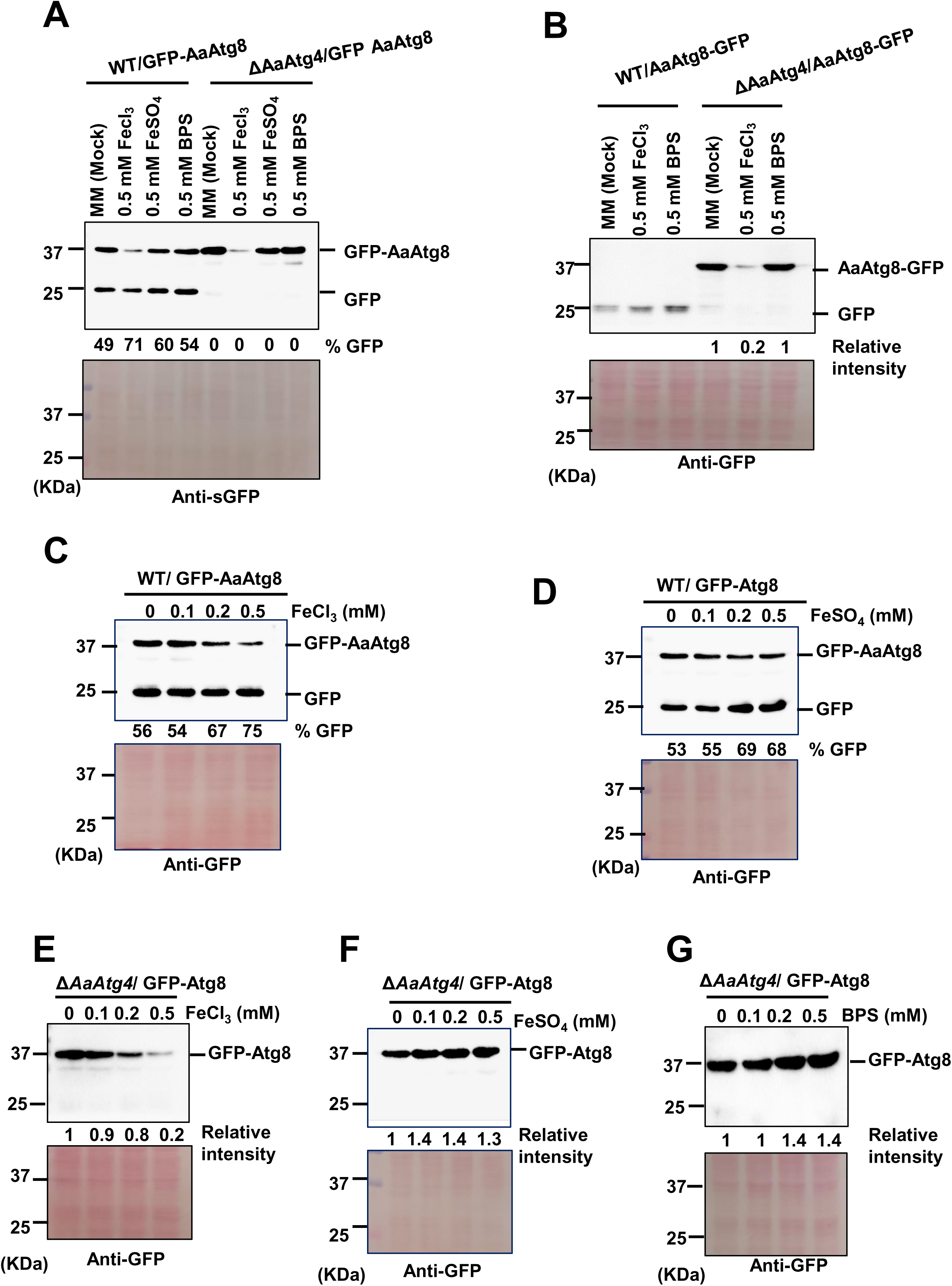
Iron modulates AaAtg8 stability without affecting autophagy or AaAtg4 activity. (A) In the AaAtg8 proteolysis assay, free GFP accumulation is comparable in the WT/GFP-AaAtg8 strain grown in minimal medium (MM) or MM supplemented with ferric chloride (FeCl_3_), ferrous sulfate (FeSO_4_), or the ferrous iron chelator bathophenanthrolinedisulfonic acid (BPS). Deletion of *AaAtg4* abolishes free GFP release under all conditions. (B) In the AaAtg4 protease assay, free GFP is fully released from AaAtg8-GFP in the wild type grown in MM or MM supplemented with FeCl_3_ or BPS, whereas no release is detected in the Δ*AaAtg4* strain. (C–D) GFP-AaAtg8 fusion protein levels are markedly reduced following FeCl_3_ treatment compared to FeSO_4_. In the wild type, GFP-AaAtg8 levels decrease progressively with increasing FeCl_3_ concentration (C), while FeSO_4_ has only a minor effect (D). (E–G) A similar FeCl_3_-dependent reduction of GFP-AaAtg8 is observed in the Δ*AaAtg4* strain (E). Conversely, GFP-AaAtg8 levels increase slightly with higher concentrations of FeSO_4_ (F) or BPS (G).

### Iron modulates AaAtg4 phosphorylation in a concentration-dependent manner

Phos-tag analysis revealed that AaAtg4 was phosphorylated in the wild-type strain cultured in MM. Upon supplementation with 0.5 mM FeCl_3_ or FeSO_4_, both AaAtg4 protein abundance and phosphorylation levels decreased (Fig. 7A). Conversely, treatment with BPS enhanced AaAtg4 abundance and phosphorylation. A dose–response assay further showed that low concentrations of FeCl_3_ (0.1–0.2 mM) promoted AaAtg4 phosphorylation. In contrast, higher concentrations (0.5 mM) suppressed it (Fig. 7B). Western blot analysis of proteins from the WT/HA-AaAtg8 strain grown in MM or MM supplemented with varying concentrations of FeCl_3_ consistently produced a single 18 kDa band (Fig. 7C), with no detectable HA-AaAtg8-PE (∼16 kDa). Again, the abundance of HA-AaAtg8 decreased as the FeCl_3_ concentration increased. Fluorescence microscopy analysis revealed that the WT/sGFPN-AaAtg8+sGFPC-AaAtg4 strain exhibited comparable green fluorescence intensity when cultured in MM or in MM supplemented with 0.5 mM FeCl₃, FeSO₄, or BPS (Fig. 7D), indicating that iron exerts no drastic effect on the interaction between AaAtg4 and AaAtg8.

**FIG 7.**
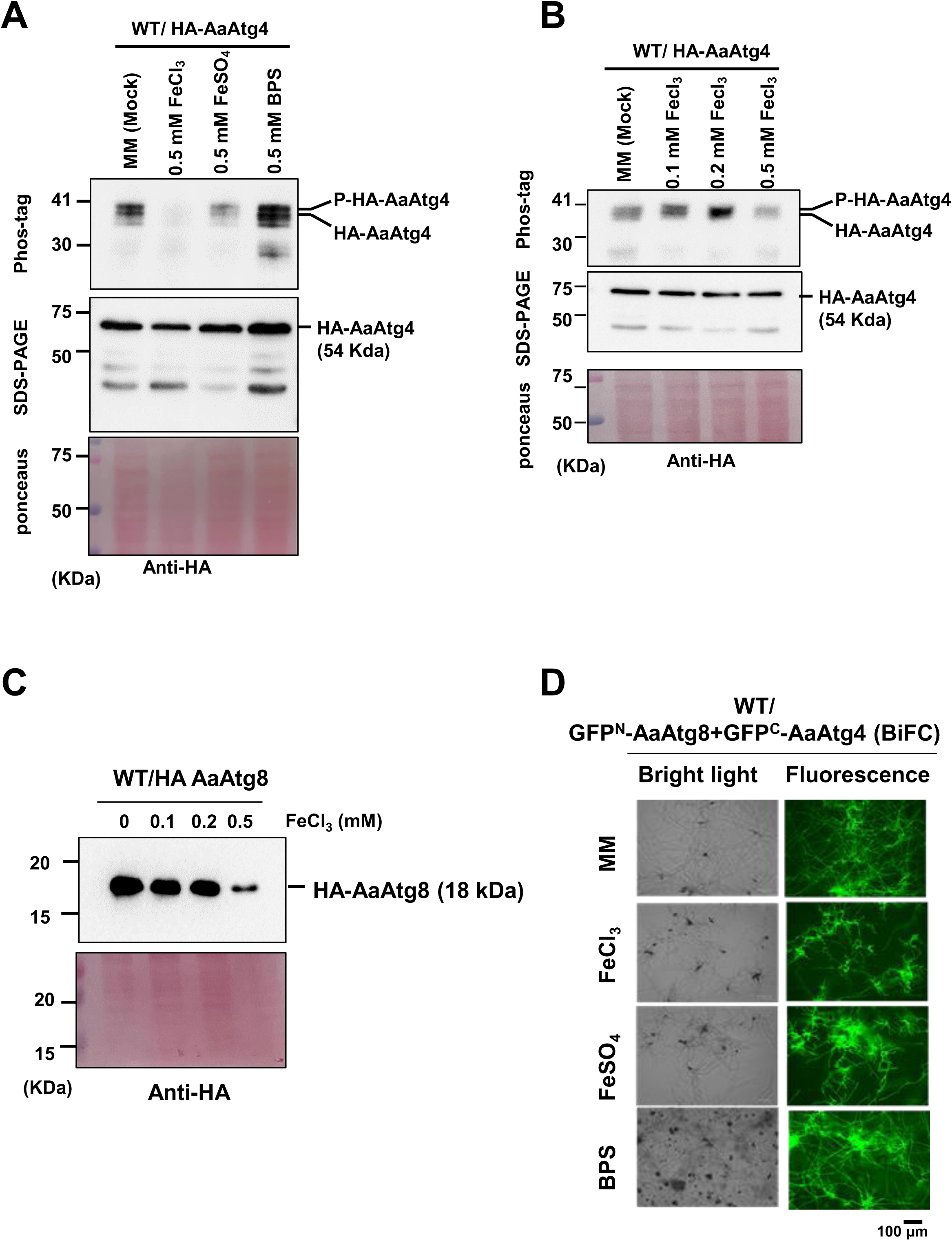
Iron modulates AaAtg4 phosphorylation in a concentration-dependent manner. (A) Phos-tag analysis shows that AaAtg4 is phosphorylated in the wild-type strain grown in minimal medium (MM). Supplementation with 0.5 mM FeCl₃ or FeSO₄ reduces both AaAtg4 protein abundance and phosphorylation, whereas treatment with the iron chelator BPS enhances them. (B) Dose–response analysis reveals that low concentrations of FeCl₃ (0.1–0.2 mM) promote AaAtg4 phosphorylation, while higher concentrations (0.5 mM) suppress it. (C) Western blotting of the WT/HA-AaAtg8 strain cultured in MM or MM supplemented with FeCl₃ shows a consistent 18 kDa band corresponding to HA-AaAtg8, with no detectable HA-AaAtg8-PE (∼16 kDa). (D) Fluorescence microscopy of the WT/sGFPN-AaAtg8+sGFPC-AaAtg4 strain reveals comparable GFP intensity in MM or MM supplemented with 0.5 mM FeCl₃, FeSO₄, or BPS, indicating that iron does not markedly affect the AaAtg4–AaAtg8 interaction.

## DISCUSSION

Autophagy is highly sensitive to redox conditions (7, 8). In this study, we investigated the effects of DTT and iron on autophagy, with a focus on AaAtg4. DTT, a strong reducing agent with two SH groups, can be oxidized to form a disulfide bond (13). Our results showed that DTT markedly enhances autophagy in *A. alternata* in a concentration-dependent manner. Moderate DTT concentrations (5-10 mM) optimally stimulate autophagic flux, whereas higher concentrations (30 mM) may exert inhibitory or cytotoxic effects, reducing efficiency. Notably, this effect is independent of AaAtg4 protease activity, which remains constitutively efficient and unaffected by DTT-induced redox changes. Thus, DTT appears to promote autophagy primarily by stimulating downstream processes such as autophagosome formation and AaAtg8 turnover, rather than by directly modulating AaAtg4 activity. Alternatively, DTT may promote autophagic flux through stress-signaling pathways, likely involving ER stress and ROS accumulation (11, 25). Consistent with findings in yeast and mammalian systems, ER stress and redox imbalance activate upstream autophagy regulators such as the unfolded protein response (UPR) and stress-responsive kinases, without directly altering Atg4 function (26, 27).

DTT modulates the interaction between AaAtg4 and AaAtg8 in *A. alternata*. At concentrations between 0.5 and 5 mM, DTT significantly enhances the fluorescence signal of the BiFC strain, indicating that reducing conditions promote or stabilize the AaAtg4–AaAtg8 interaction. This enhanced interaction likely reflects increased efficiency of AaAtg8 processing, which is essential for autophagosome formation. However, when cultured in 10 mM DTT, fluorescence intensity drops sharply, suggesting that excessive reduction disrupts protein folding or cellular homeostasis, thereby impairing the AaAtg4–AaAtg8 interaction. High concentrations of DTT are known to induce reductive stress, destabilizing protein structures and interfering with normal cellular processes (28, 29).

DTT promotes AaAtg8 lipidation in a concentration-dependent manner (Fig. 4A), suggesting that reducing conditions favour the conjugation of AaAtg8 to PE, a critical step in autophagosome membrane expansion. This effect may arise because DTT maintains the cysteine residues of AaAtg4 in a reduced state, which is necessary for processing AaAtg8 prior to lipidation (17). However, co-treatment with H₂O₂ abolishes the formation of HA-AaAtg8-PE, even in the presence of DTT (Fig. 4B). Since H₂O₂ alone (≤20 mM) enhances AaAtg8 lipidation (16), oxidative stress itself is unlikely to interfere with the lipidation process. Instead, the antagonism appears to result from chemical interactions between DTT and H₂O₂. H₂O₂ can oxidize the thiol groups (–SH) in DTT to disulfides (–S–S–), thereby inactivating its reducing capacity [30]. Moreover, their reaction may generate reactive sulfur species (e.g., sulfenic acids) and oxygen radicals, which could destabilize protein structures or disrupt enzyme–substrate interactions (31, 32). Thus, mixing DTT and H₂O₂ neutralizes their individual effects, preventing AaAtg8 lipidation. Collectively, these findings highlight that AaAtg8 lipidation is highly sensitive to redox balance.

H₂O₂ (20–30 mM) slightly suppresses AaAtg4 activity (16), an effect that can be counteracted by DTT, consistent with its role as a reducing agent. The interaction between AaAtg4 and AaAtg8, the lipidation of AaAtg8, and the protease activity of AaAtg4 are all modulated by the cellular redox environment to varying extents. Under moderately reducing conditions induced by DTT, AaAtg4–AaAtg8 interaction is consistently enhanced, AaAtg8 lipidation is promoted, and AaAtg4 protease activity is maintained.

DTT (5 and 10 mM) or H₂O₂ (10 mM) alone enhances autophagy, whereas their simultaneous presence maintains autophagy at a moderate yet steady level (Fig. 4D). This suggests that the interplay between reductive stress (DTT) and oxidative stress (H₂O₂) establishes a cellular environment that restrains autophagic activity. Notably, increasing H₂O₂ concentrations (20–30 mM) does not proportionally elevate autophagy, indicating that the response reaches a threshold under combined stress conditions. These findings reveal that autophagy is not linearly dependent on oxidative stress intensity when reductive stress is present, but instead stabilizes at a moderate level in *A. alternata*. The coexistence of DTT and H₂O₂ may reflect a balance between redox signaling and stress adaptation, producing a redox state that sustains autophagy without driving cells toward excessive degradation or apoptosis (33). This steady-state autophagy likely represents a protective adaptation that preserves cellular homeostasis under dual-stress conditions (34). Overall, our results indicate that autophagy is not simply triggered by heightened oxidative or reductive stress, but rather emerges from the dynamic equilibrium between these opposing forces.

DTT enhances AaAtg4 protein abundance and phosphorylation in a concentration-dependent manner (Fig. 3E), with maximal induction observed at moderate levels (≤5 mM). At these concentrations, DTT promotes both AaAtg4 accumulation and phosphorylation, indicating that reductive stress not only increases protein stability or expression but also activates its regulatory modification, thereby facilitating autophagic flux. This effect may occur through stress-responsive transcriptional or post-translational mechanisms (11, 35). Phosphorylation of Atg4 is known to regulate its protease activity and its interaction with Atg8, thereby modulating autophagosome formation in yeast (11, 17). In *A. alternata*, the optimal DTT concentration required for AaAtg4 phosphorylation coincides with that required for autophagy induction. Beyond 5 mM, both AaAtg4 protein abundance and phosphorylation decline, suggesting that excessive reductive stress disrupts AaAtg4 regulation. This impairment may result from protein destabilization, inhibition of kinase activity, or activation of compensatory feedback mechanisms that limit autophagy (11).

Previously, we demonstrated that AaAtg4 directly interacts with AaHog1 (16), a mitogen-activated protein kinase involved in resistance to osmotic stress, fungicides, and xenobiotics (24). AaHog1 is strongly phosphorylated in response to H₂O₂. In an AaHog1-deficient mutant, AaAtg4 becomes hyperphosphorylated under conditions unfavorable for autophagy (e.g., absence of H₂O₂ or exposure to excessive H₂O₂), suggesting that AaHog1 functions as a negative regulator of AaAtg4 phosphorylation (16). While low-to-moderate concentrations of DTT increase AaAtg4 phosphorylation, likely through redox-sensitive pathways, DTT does not affect AaHog1 phosphorylation (Fig. 4F). These findings indicate that AaHog1 responds specifically to oxidative stress rather than to general redox changes and that DTT-mediated autophagy occurs independently of AaHog1 signaling, in contrast to observations in yeast (36, 37). Moreover, AaHog1 is not involved in autophagy as deleting the gene resulted in a mutant that exhibits normal autophagy under starvation and in the presence of H_2_O_2_ (data not shown). The precise biological role of the interaction between AaAtg4 and AaHog1 remains to be elucidated. It is plausible that, under severe oxidative stress, AaHog1 suppresses AaAtg4 phosphorylation, thereby restricting autophagic flux, despite not directly influencing autophagy itself. Such regulation may represent a protective adaptation, preventing excessive autophagy that could otherwise lead to detrimental self-digestion under extreme oxidative stress (38, 39).

Using the WT/GFPN-AaAtg8+GFPC-AaAtg4 (BiFC) strain, we found that the redox environment strongly influences the subcellular localization of the AaAtg4–AaAtg8 complex (Fig. 5). In the absence of DTT, green fluorescence is restricted to the hyphae and excluded from vacuoles. Upon DTT treatment, fluorescence is uniformly distributed throughout the hyphae, including vacuoles, indicating that reducing conditions promote vacuolar localization of the AaAtg4–AaAtg8 complex. This effect may reflect enhanced autophagic flux or stabilization of protein–protein interactions under reductive stress (39, 40). In contrast, high concentrations of H₂O₂ (40 mM) abolish the DTT-induced vacuolar localization, suggesting that oxidative stress disrupts vacuolar targeting, potentially by oxidizing cysteine residues in AaAtg4 or impairing AaAtg8 membrane conjugation. Notably, only strong oxidative stress overrides the reducing environment established by DTT, as lower H₂O₂ concentrations (≤30 mM) do not produce this effect.

We previously showed that autophagy influences siderophore biosynthesis, iron acquisition, and homeostasis in *A. alternata* (15, 21). In this study, we further examined the role of iron, a known reducing agent. Iron modulates autophagic flux, without affecting AaAtg4 activity, AaAtg8 lipidation, or AaAtg4–AaAtg8 interactions. Instead, iron modulates AaAtg4 phosphorylation in a concentration-dependent manner (Fig. 7A, B): low levels of FeCl₃ enhance phosphorylation, whereas higher levels suppress it, accompanied by reduced protein abundance. This biphasic regulation suggests that iron functions as a signaling factor (41), modulating AaAtg4 activity through post-translational modification. In contrast, treatment with the iron chelator BPS increases both AaAtg4 phosphorylation and protein abundance. Interestingly, iron supplementation differentially affects AaAtg8 stability (Fig. 6), independent of AaAtg4-mediated proteolysis. FeCl₃ markedly reduces GFP-AaAtg8 and AaAtg8-GFP fusion protein levels in a concentration-dependent manner, whereas FeSO₄ exerts only minor effects.

Conversely, chelation of ferrous iron by BPS slightly increases AaAtg8 abundance, possibly by altering protein turnover or folding (42). The distinction between ferric and ferrous iron highlights the importance of oxidation state (Fe^2+^ vs. Fe^3+^) in determining the stability of AaAtg8. Moreover, iron acts as a molecular switch for AaAtg4 phosphorylation, suggesting that *A. alternata* can sense iron availability and improve autophagy-related processes accordingly. Although iron overload can induce ferroptosis, an iron-induced cell death characterized by lipid peroxidation and membrane damage (10), *A. alternata* can tolerate high iron levels without exhibiting lipid peroxidation or cell death, partly due to autophagy-mediated iron detoxification. Autophagy impairment leads to the intracellular accumulation of Fe^2+^ in *A. alternata* (21).

Nonetheless, iron exerts a dual regulatory role: it destabilizes AaAtg8 protein abundance and modulates AaAtg4 phosphorylation, thereby increasing autophagic flux. This selective regulation may represent an adaptive strategy that balances the stability of autophagy-related proteins with environmental iron levels, thereby optimizing cellular responses to nutrient stress.

## CONCLUSIONS

Autophagy in *A. alternata* is tightly regulated by redox balance and iron availability. AaAtg4 plays a critical role in the lipidation and delipidation of AaAtg8 during autophagy. As summarized in Fig. 8, low-to-moderate reductive stress induced by DTT promotes AaAtg4–AaAtg8 interactions and AaAtg4 phosphorylation, whereas excessive reduction destabilizes these processes. However, increasing DTT concentrations proportionally elevate AaAtg8 lipidation and autophagy, whereas DTT has no effect on AaAtg4 protease activity. Previous studies have demonstrated that oxidative stress via H₂O₂ stimulates autophagy at moderate levels but suppresses AaAtg4 activity and vacuolar localization under severe conditions. In this study, we found that the interplay between reductive and oxidative stress sustains autophagy at a protective steady state. Iron modulates AaAtg4 phosphorylation and AaAtg8 stability, thereby contributing to protein homeostasis and autophagic flux. Collectively, these findings reveal that autophagy is governed by a balanced regulatory network integrating redox signals and nutrient cues, enabling adaptive responses to environmental stress.

**FIG 8.**
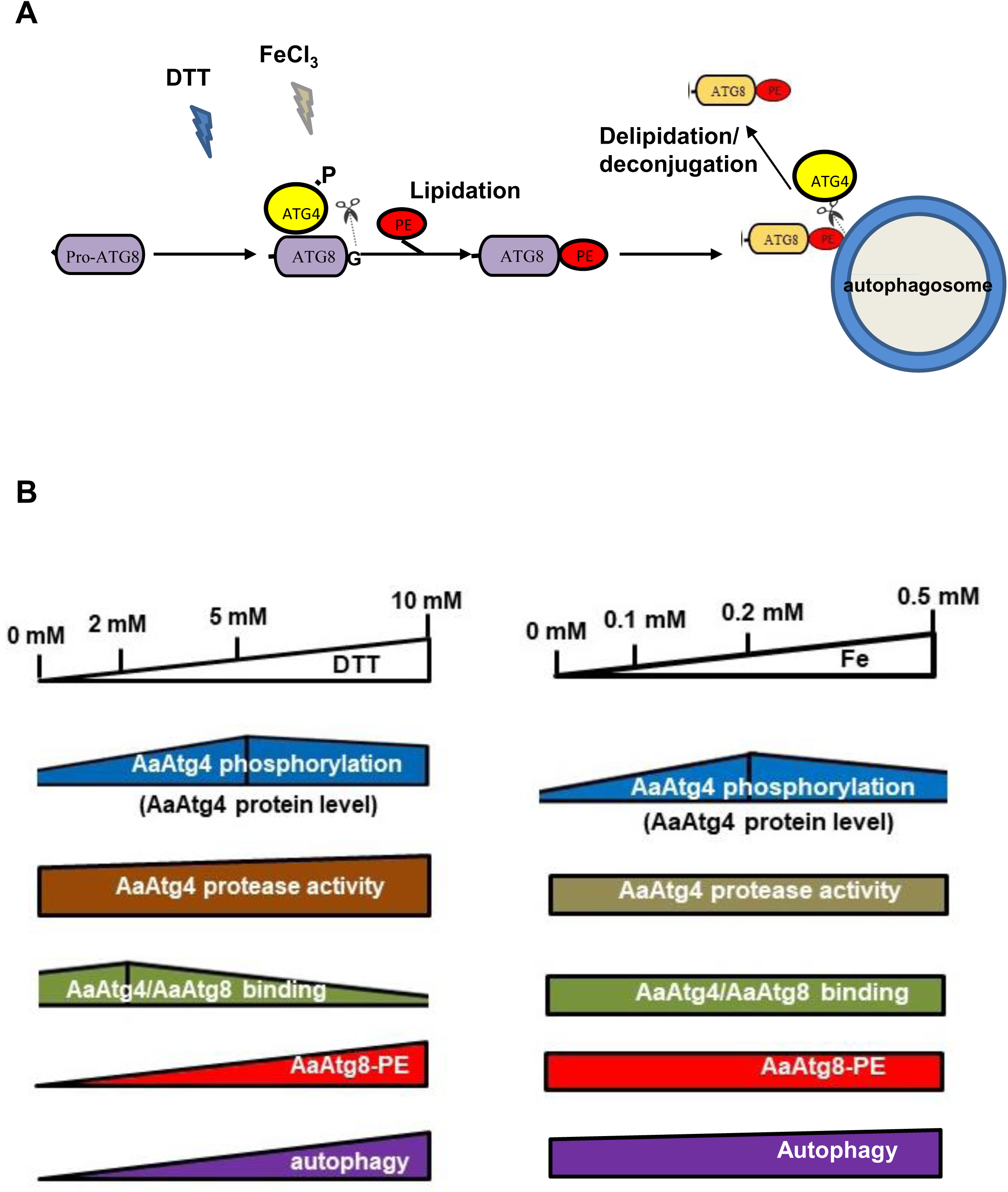
DTT and iron regulate autophagy via modifying AaAtg4 protease. (A) A schematic illustrates the regulation of autophagy through AaAtg4 protease in response to environmental iron and DTT. AaAtg4 cleaves arginine from the C-terminus of AaAtg8, exposing its glycine residue, which is subsequently conjugated to phosphatidylethanolamine (PE) to form the Atg8-PE complex, a process known as lipidation. DTT also promotes the AaAtg4–AaAtg8 interaction. In addition to catalyzing lipidation, AaAtg4 also mediates delipidation, releasing AaAtg8 from Atg8-PE anchored on the outer membrane of autophagosomes. (B) Comparative analyses of AaAtg4 activity under varying concentrations of DTT or FeCl₃ reveal that DTT enhances AaAtg8 lipidation and autophagic flux. Furthermore, low levels of DTT promote AaAtg4 phosphorylation and strengthen the AaAtg4–AaAtg8 interaction. In contrast, iron does not influence autophagic flux, AaAtg4 activity, AaAtg8 lipidation, or AaAtg4–AaAtg8 interactions. At concentrations of 0.1–0.2 mM, iron enhances AaAtg4 phosphorylation, whereas at 0.5 mM it suppresses this modification. Neither DTT nor iron affects AaHog1 phosphorylation, even though AaAtg4 interacts with AaHog1.

## MATERIALS AND METHODS

### Fungal strains and culture conditions

The wild-type EV-MIL31 strain of *A. alternata* (Fr.) Keissler used in this study was isolated from a diseased Minneola tangelo (*Citrus paradisi* Macfad. x *C. reticulata* Blanco) leaf (43). The Δ*AaAtg4* strain was previously identified in EV-MIL31 by deleting the AaAtg4 gene encoding a protease using a split-marker approach (22). Fungal strains used in this study were obtained from Lu et al. (16), and their characteristics and intended applications are summarized in Supplementary Table S1. Unless otherwise stated, fungi were grown on potato dextrose agar (PDA, Difco, Sparks, MD, U.S.A.) or minimum medium (MM) (44) under constant fluorescent light at 28 °C for 3 to 5 days. For protein isolation, fungal strains were cultured in PDB (Difco) for 24 h. A medium shift was performed in all experiments. Fungal strains were cultured in PDB for 24 h, after which the mycelium was harvested by filtration through Miracloth, rinsed with sterile water, and transferred to PDB or MM. To assess the effect of iron, the mycelium was transferred to MM or MM supplemented with varying concentrations of FeCl₃, FeSO₄, or BPS (a ferrous iron chelator), and incubated for 4 h. To examine the effect of DTT, fungal mycelium was transferred to PDB or PDB supplemented with different concentrations of DTT. To determine whether DTT counteracts the effect of H₂O₂, the test strain was transferred to either PDB (one flask) or PDB supplemented with 30 mM H₂O₂ (two flasks) and incubated for 4 h. DTT was subsequently added to one of the H₂O₂-treated cultures, followed by an additional 2 h of incubation. Mycelium was then harvested and subjected to protein extraction and western blot analysis.

### Genetic modification in *A. alternata*

For the proteolysis assay, a GFP-AaAtg8 fusion fragment was constructed by joining *AaAtg8* cDNA with the C terminus of GFP coding sequence and expressed under the control of the *AaAtg8* promoter and cloned into a pCB1532 plasmid, carrying a sulfonylurea-resistance (Sur) gene. The resulting plasmid was transformed into protoplasts prepared from the wild-type (WT) and the Δ*AaAtg4* strains to yield WT/GFP-AaAtg8 and Δ*AaAtg4*/GFP-AaAtg8 strains, respectively. For the AaAtg4 protease activity assay, an AaAtg8-GFP fusion fragment was made by fusing AaAtg8 cDNA with the N terminus of GFP and cloned into pCB1532, and the resulting plasmid was transformed into WT and Δ*AaAtg4* to yield WT/AaAtg8-GFP and Δ*AaAtg4*/AaAtg8-GFP strains, respectively. *AaAtg4* and *AaAtg8* were cloned into a pNeoGPE1-HA plasmid carrying a linker HA-tag sequence (5’-atgggcagctacccatacgatgttccagattacgct-3’), and a neomycin (G418)-resistant cassette, and the resulting construct was transformed into wild-type protoplasts to yield WT/HA-AaAtg4 and WT/HA-AaAtg8 strains, respectively. The fungal strain used for bimolecular fluorescence complementation (BiFC) assay was created as follows. The N-terminal half of GFP (GFP^N^) was fused to AaAtg8 and cloned into pHygroGPE1, carrying a hygromycin-resistant cassette, to yield a GFP^N^-AaAtg8 plasmid. The C-terminal half of GFP (GFP^C^) was fused to AaAtg4 and cloned into pNeoGPE1 to yield a GFP^C^-AaAtg4 plasmid. Both GFP^N^-AaAtg8 and GFP^C^-AaAtg4 were co-transformed into protoplasts prepared from WT to yield a WT/GFP^N^-AaAtg8+GFP^C^-AaAtg4 strain.

### Western blot and phos-tag analyses

Proteins were isolated, separated by SDS-PAGE, and transferred to polyvinylidene difluoride (PVDF) membranes as described (15). Proteins were quantified using the Bradford assay (Bio-Rad, Hercules, CA, U.S.A.) and stained with Ponceau S after electrophoresis for loading control. Proteins on PVDF were incubated with a rabbit anti-GFP antibody (1:5000) overnight at 4 °C and reacted with an HRP-conjugated goat anti-rabbit IgG (1:10,000, Jackson ImmunoResearch, West Grove, PA, U.S.A.) and LumiFlash Prime Chemiluminescent Substrate, HRP (Visual Protein, Taipei, Taiwan). Chemiluminescence signals were acquired using the ChemiDoc MP imaging system (Bio-Rad) and ImageLab software (version 6.1.0). Band intensity was determined using ImageJ software (US National Institutes of Health, Bethesda, MD, U.S.A.) (https://imagej.net/ij/). The percentage of free GFP released from GFP–AaAtg8 was calculated by dividing the intensity of the free GFP band by the total intensity of free GFP plus GFP–AaAtg8, and then multiplying by 100%. For phos-tag assays, proteins were isolated from fungi using a lysis buffer supplemented with protease inhibitors, as described by Lu et al. (16). Proteins were mixed with 1 mM Zn (NO_3_)_2_ at a 10:1 (v/v) ratio and 5x loading dye before loading. To prepare Zn-Phos-tag-SDS-PAGE gels, 10 mM Zn (NO_3_)_2_ and 5 mM Phos-tag acrylamide AAL-107 (Fujifilm Wako, Chuo-Ku, Osaka, Japan) were added to a 30% acrylamide:bis-acrylamide ( 37.5:1, v/v) solution. After electrophoresis, proteins were transferred to a PVDF membrane, rinsed with 10 mM EDTA, and incubated with an anti-HA antibody (Sigma-Aldrich) and a secondary antibody (Santa Cruz Biotechnology, Dallas, TX, U.S.A.).

### AaAtg8 lipidation assay

The WT/HA-AaAtg8 strain was used to determine the attachment of PE to AaAtg8, a process known as lipidation (4), during autophagy. The strain was grown in PDB for 24 h, then shifted to a newly prepared PDB supplemented with different concentrations of DTT and incubated for 2-4 h. Proteins were extracted and separated by SDS-PAGE containing 6 M urea as described by Hirata et al. (45), and HA-AaAtg8-PE was detected by Western blots using an anti-HA antibody as described above. The HA-AaAtg8-PE protein migrated faster than the HA-AaAtg8 in SDS-PAGE gel due to its higher hydrophobicity (23).

### Fluorescence microscopy

Green fluorescence was observed using a ZOE Fluorescent Cell Imager (Bio-Rad) microscope equipped with a fluorescence filter set with excitation at 488 nm and emission at 510 nm. Vacuoles were stained with CMAC (7-amino-4-chloromethylcoumarin, 100 μM; Thermo Fisher Scientific, Waltham, MA, U.S.A.) and visualized with excitation at 353 nm and emission at 466 nm.

## DATA AVAILABILITY

All data supporting this study are available in the Figshare repository https://figshare.com/account/items/32074998/edit; doi: https://10.6084/m9.figshare.32074998. Gene sequences are available in the NCBI database under accession numbers: KAH8643735 (AaAtg4) and OK617334 (AaAtg8).

